# Spatial Regulation of Polo-like Kinase Activity during *C. elegans* Meiosis by the Nucleoplasmic HAL-2/HAL-3 Complex

**DOI:** 10.1101/534339

**Authors:** Baptiste Roelens, Consuelo Barroso, Alex Montoya, Pedro Cutillas, Weibin Zhang, Alexander Woglar, Chloe Girard, Enrique Martinez-Perez, Anne M. Villeneuve

## Abstract

Proper partitioning of homologous chromosomes during meiosis relies on the coordinated execution of multiple interconnected events: Homologs must locate, recognize and align with their correct pairing partners. Further, homolog pairing must be coupled to assembly of the synaptonemal complex (SC), a meiosis-specific tripartite structure that maintains stable associations between the axes of aligned homologs and regulates formation of crossovers between their DNA molecules to create linkages that enable their segregation. Here we identify HAL-3 (<u>H</u>omolog <u>A</u>lignment 3) as an important player in coordinating these key events during *C. elegans* meiosis. HAL-3 and the previously-identified HAL-2 are interacting and interdependent components of a protein complex that localizes to the nucleoplasm of germ cells. *hal-3* (or *hal-2*) mutants exhibit multiple meiotic prophase defects including failure to establish homolog pairing, inappropriate loading of SC subunits onto unpaired chromosome axes, and premature loss of synapsis checkpoint protein PCH-2. Further, loss of *hal* function results in misregulation of the subcellular localization and activity of polo-like kinases (PLK-1 and PLK-2), which dynamically localize to different defined subnuclear sites during wild-type prophase progression to regulate distinct cellular events. Moreover, loss of PLK-2 activity partially restores tripartite SC structure in a *hal* mutant background, suggesting that the defect in pairwise SC assembly in *hal* mutants reflects inappropriate PLK activity. Together our data support a model in which the nucleoplasmic HAL-2/HAL-3 protein complex constrains both localization and activity of meiotic Polo-like kinases, thereby preventing premature interaction with stage-inappropriate targets.

## Introduction

Generation of haploid gametes during sexual reproduction depends on the ability of homologs to form physical connections that will enable them to accurately orient and be partitioned during the first meiotic division. To form these links, homologous chromosomes undergo an extended prophase program during which they must locate, pair and align with their correct pairing partner to form recombination-based interactions that will mature into these physical connections. Importantly, these events are not independent and need to be tightly coordinated to ensure reproductive success. For instance, pairing between homologous chromosomes is stabilized by a tripartite structure, the synaptonemal complex (SC), that assembles at the interface between the two homologs, a process known as synapsis. Although the SC normally assembles only between paired homologous segments, the structure itself is indifferent to homology, and therefore pairing needs to be coordinated with synapsis to ensure that only fruitful homologous interactions are stabilized.

Coordination of the events in the model organism *C. elegans* is achieved in part through the regulated action of protein kinases. Upon entry into meiotic prophase, CHK-2 phosphorylates a family of related Zinc-finger DNA binding proteins (HIM-8, ZIM-1/2/3), which bind preferentially to chromosomal sites known as Pairing Centers (PCs) that are located near one end of each chromosome. PCs mediate connections between the chromosomes and the cytoplasmic motility apparatus through association with protein complexes that span the nuclear envelope (NE) (MACQUEEN *et al.* 2005; PHILLIPS *et al.* 2005; PHILLIPS AND DERNBURG 2006; PENKNER *et al.* 2007; PENKNER *et al.* 2009; SATO *et al.* 2009). CHK-2-mediated phosphorylation promotes accumulation of the PC-binding proteins at their respective PCs and also creates a phospho-motif that serves as a docking site for the Polo Box Domain of the Polo-like kinases PLK-2 and PLK-1. This localized PLK activity at the PC triggers and sustains the NE association and chromosome movements that promote selective stabilization of pairing and assembly of the SC between homologs (HARPER *et al.* 2011; LABELLA *et al.* 2011; KIM *et al.* 2015). Once homologs are paired and synapsed, the meiotic prophase specific Polo-like kinase PLK-2 is released from the PC – NE attachment sites and becomes associated with the SCs, initially localizing along their full length and eventually becoming restricted to limited subdomains of the SCs (MACHOVINA *et al.* 2016; NADARAJAN *et al.* 2017; PATTABIRAMAN *et al.* 2017; FERRANDIZ *et al.* 2018). This sequential association of PLK-2 with distinct nuclear subdomains during prophase progression appears to be important to ensure the proper execution of the prophase program. For example, a *him-8* mutation that prevents association of PLK-2 to the X chromosome PCs leads to defective pairing and synapsis of the X chromosomes (PHILLIPS *et al.* 2005; HARPER *et al.* 2011). Similarly, PLK-1/2-dependent phosphorylation of substrates such as the SC-CR (SC central region) component SYP-4 or the axis component HTP-1 have been shown to be important, respectively, for limiting initiation of meiotic recombination and for protection of sister-chromatid cohesion during the first meiotic division (NADARAJAN *et al.* 2017; FERRANDIZ *et al.* 2018). How the dynamic localization of PLK-2 contributes to ensuring proper coordination of the different prophase events, and the mechanisms responsible for the spatio-temporal regulation of PLK activity, are still poorly understood.

We provide here new insights into the coordination of meiotic prophase through analysis of the functions of HAL-2/HAL-3, a protein complex that is concentrated in germ cell nuclei. We identify HAL-3 as an interacting partner of HAL-2, which was previously described as an important regulator of meiotic prophase events (ZHANG *et al.* 2012), and we show that these two factors are interdependent for their localization in the nucleoplasm of germ cells. Loss of *hal-3* function induces defects similar to the *hal-2* mutant, including defective homolog pairing, lack of markers of PC-led chromosome movements, and inappropriate loading of components of the central region of the SC onto unpaired homologs. Importantly, we also show that in the absence of either HAL-2 or HAL-3, Polo-like kinases are mis-localized and active on untimely substrates. Further, removal of the major meiotic prophase Polo-like kinase PLK-2 is sufficient to rescue some of the meiotic defects observed in the *hal-2* mutant background. Our analysis suggests that the HAL-2/HAL-3 complex provides a previously unappreciated layer of regulation of meiotic progression, by preventing premature interaction of meiotic Polo-like kinases with stage-inappropriate targets. Further, our data suggest that HAL-2/HAL3 accomplishes this not only by promoting activity of CHK-2 but also by antagonizing the activity of PLK-2.

## Materials and Methods

### Strains and genetics

Unless otherwise specified, worms were cultured at 20°C under standard conditions (BRENNER 1974). The following *C. elegans* strains were used:

- N2
- ATGSi43: *fqSi37[htp-1::gfp Cb-unc-119(+)] II; htp-1(gk174)* IV
- AV158: + / *nT1 IV; rad-50(ok197) / nT1[unc-?(n754) let-? qIs50]* V
- AV327: *hal-2(me79)/qC1[qIs26]* III
- AV659: *hal-2(me79) III; meIs10[pie-1p::gfp::hal-2 Cb-unc-119(+)]*
- AV660: *chk-2(me64) rol-9(sc148)/ sC4 [dpy-21(e428)] V*
- AV684: + / *hT2[qIs48] I; hal-2(me79) / hT2* III
- AV691: + / *hT2[qIs48] I; hal-2(me79) / hT2 III; chk-2(me64) rol-9 (sc148) / + V*
- AV706: *meIs10[pie-1p::gfp::hal-2 Cb-unc-119(+)]*
- AV799: *hal-3(me99) / mIn1[dpy-10(e128) mIs14]* II
- AV897: *hal-3(me99) / mIn1[dpy-10(e128) mIs14]* II; + / *nT1 IV; rad-50(ok197) / nT1[unc-?(n754) let-? qIs50]* V
- AV928: *plk-2(ok1936) / hT2[qIs48] I; hal-2(me79) / hT2* III
- AV932: *ieDf2 / nT1 IV; + / nT1[unc-?(n754) let-? qIs50]* V
- AV933: *hal-3(me99) / mIn1[dpy-10(e128) mIs14]* II; *ieDf2 / nT1* IV; + / *nT1[unc-?(n754) let-? qIs50]* V
- AV934: *meSi3[hal-3p::hal-3::gfp::hal-3 3’UTR Cb-unc-119(+)] II; unc-119(ed3)* III
- AV984: *fqSi37[htp-1::gfp Cb-unc-119(+)] II; hal-2(me79)/qC1[qIs26] III; htp-1(gk174)* IV
- AV1034: + / *hT2[qIs48] I; meSi3[hal-3p::hal-3::gfp::hal-3 3’UTR Cb-unc-119(+)]* II; *hal-2(me79) / hT2* III
- CA896: *plk-2(ok1936) / hT2[qIs48]* I

### Immunoprecipitation of GFP::HAL-2

Nuclear purification and preparation of nuclear soluble and insoluble fractions were performed as described in (SILVA *et al.* 2014). Approximately 400 μg of total protein for each nuclear fraction was incubated with 50 μl of GFP-TRAP agarose beads for 2 h and 30 min at 4 °C. The beads were then washed at least 5 times and collected by centrifugation at 2700xg for 2 min at 4 °C. Each sample was resuspended in 30μl of SDS-sample buffer and protein complexes were eluted by boiling.

### Mass spectrometry

Immunoprecipitated proteins from the different fractions (nuclear soluble and insoluble) were run on a 10% acrylamide gel for 20 minutes (fixed 80V), and then stained overnight using Coomassie brilliant blue. Each gel lane was excised into 4 equally sized horizontal bands along the entire length of the gel lane. Gel pieces were subjected to overnight trypsin digestion at 37 °C using an adaptation of the gel digestion procedure of (SHEVCHENKO *et al.* 2006). Peptides were recovered post-digestion and subsequently dried to completeness using a vacuum centrifuge. Dried peptides were resuspended in 0.1% trifluoroacetic acid (TFA) and subjected to LC-MS analysis using an Ultimate 3000 nano HPLC coupled to a LTQ-Orbitrap-XL mass spectrometer. Peptides were loaded onto a trap column (Acclaim PepMap 100 C18, 100μm × 2cm) at 8μL/min in 2% acetonitrile, 0.1% TFA. Peptides were then eluted on-line to an analytical column (Acclaim Pepmap RSLC C18, 75μm × 50cm). A 90-minute gradient separation was used, 4-35% of buffer B (composition: 80% acetonitrile, 0.1% formic acid). Eluted peptides were analysed by the LTQ-XL operating in positive polarity using a data-dependent acquisition mode. A top 7 CID method was implemented with full MS scans acquired in the Orbitrap mass analyzer (range m/z 350–1500) with a mass resolution of 15,000 (at m/z 400). The 7 most intense peaks with charge state ≥2 were fragmented in the ion trap with a normalized collision energy of 35%, activation Q = 0.25 and activation time of 30 milliseconds. Maximum allowed ion accumulation times were 500 ms for full scans and 100 ms for CID. Data were processed using the MaxQuant software platform (v1.5.3.8) (COX *et al.* 2014), with database searches carried out by the in-built Andromeda search engine against the TrEMBLE C. elegans database (Downloaded – 15th October 2015).

A reverse decoy database approach was used at a 1% FDR for peptide spectrum matches and protein identification. Search parameters included: maximum missed cleavages set to 2, fixed modification of cysteine carbamidomethylation and variable modifications of methionine oxidation and protein N-terminal acetylation. Label-free quantification was enabled with an LFQ minimum ratio count of 2.

### Western blotting

Worms were collected and washed in M9 before being resuspended in Laemmli Sample Buffer (BioRad) containing .2M Dithiothreitol (DTT). Worm extracts were then boiled and separated on 4-15% SDS-PAGE gradient gel (50 worms per lane). Western analysis was performed using standard procedures with rabbit anti-HAL-2 antibody (1:10,000) (ZHANG *et al.* 2012).

### RNA depletion

RNAi experiments were performed by placing worms at 15°C on NGM plates containing ampicillin (100μg/mL) and IPTG (1mM), seeded with bacteria carrying the appropriate clone from the Ahringer library (KAMATH AND AHRINGER 2003) for depletion of *rad-50*. Depletion of *hal-3*(*T15H9.2*) was performed under similar experimental conditions using the plasmid pBR7, which was obtained by blunt end cloning of a PCR product obtained by amplification of genomic DNA extracted from the wild-type N2 using the primers oBR224 (ccctgagccgtgtaagttgt) and oBR225 (cccaattttccgaaagagg) into an EcoRV-digested L4440 vector (TIMMONS AND FIRE 1998).

### Transgenesis

The transgene allowing expression of the HAL-3::GFP fusion protein was obtained using the Mos Single Copy Insertion strategy (MosSCI, (FROKJAER-JENSEN *et al.* 2008)), using the *ttTi5605* insertion on chromosome *II* as a landing site. The donor plasmid, pBR224, was obtained by assembling fragments carrying the upstream, coding and downstream genomic sequences of *hal-3* together with a fragment containing a version of GFP optimized for germline expression (FROKJAER-JENSEN *et al.* 2016), into pBR49, a derivative of pCFJ350 modified for to enable type IIs restriction/ligation cloning (ENGLER *et al.* 2008). The genomic fragments were obtained by PCR amplification of wild-type genomic DNA using the following primer pairs: oBR727 (cgtcgatgcacaatccGGTCTCaCCTGgtattttcattgtcacccaaaaattg) and oBR728 (agtggaatgtcagGGTCTCaCATAttctgaaatatttttttacatgaaacaaaag) for the upstream sequence, oBR408 (cgatgcacaatccGGTCTCaTATGGCGGCAGCAAAAAGAAAATTAG) and oBR409 (agtggaatgtcagGGTCTCaCTCCTTTTTGTGCCTGACGAATTCGATC) for the coding sequence, oBR708 (cgtcgatgcacaatccGCTCTTCaTAAaatttttttttttttcgttttgccctg) and oBR709 (agtggaatgtcagGCTCTTCaGTCAcatcattgtaggtgatttcagtgg) for the downstream sequence. The GFP fragment was obtained by PCR amplification of pCFJ1848 (FROKJAER-JENSEN *et al.* 2016), using oBR406 (cgatgcacaatccGGTCTCaGGAGGTGGATCATCCTCCACATCATCCT) and oBR407 (agtggaatgtcagGGTCTCaTTTATGGGGAAGTACCGGATGACG). Correct assembly of all fragments within the donor plasmids was verified by sequencing.

### Generation of a hal-3 mutant

The *hal-3* mutant allele, *me99*, was obtained by injection of pBR1, a derivative of pDD162 (DICKINSON *et al.* 2013), allowing expression of the Cas9 nuclease and a single guide RNA (sgRNA) targeting the sequence GTCTCCCAATTTTCCGAAAGAGg, located at the end of the first exon of the *hal-3* coding sequence, into wild-type N2 worms. Individual descendants of injected hermaphrodites were selected for the expression of a co-injection marker (*Pmyo-3::mCherry*, pCFJ104) and status of the *hal-3* locus in these isolates was analyzed by PCR using primers oBR73(actgaatatcatactccaaaa), oBR74(ctcccaattttccgaaag) and oBR75(tgtaatccttttgtttcatgt) and confirmed by sequencing of the PCR fragment obtained using primers oBR73 and oBR74. The *me99* allele isolated using this strategy contains a one base pair deletion at the expected cut site of the Cas9/sgRNA complex: GTCTCCCAATTTTCCG-AAGAGg.

### Yeast two-hybrid

Yeast two-hybrid experiments were performed as described in (GIRARD *et al.* 2018). In short, cDNA sequences of *hal-2* and *hal-3* were amplified from Trisol extracted, reverse-transcribed RNAs from N2 worms using primers containing SpeI and AvrII restriction sites to allow for cloning into pDP133 (prey vector) and pDP135 (bait vector), respectively (KRAFT *et al.* 2012). Pairs of prey and bait plasmids were transformed into yeast strain YCK580 and spotted on selective media lacking both Leucine and Tryptophan (-L-W) or Leucine, Tryptophan and Histidine (-L-W-H). The primer pairs used for amplification of cDNA sequences were oBR714 (CTAACTAGTATGAACACACCGGTAAAAGCAAAA) and oBR715 (CTACCTAGGATTGAAGAATTCGTCGAATTCGTTTT) for *hal-2*, and oBR716 (CTAACTAGTATGGCGGCAGCAAAAAGAAAATTA) and oBR717 (CTACCTAGGTTTTTGTGCCTGACGAATTCGATC) for *hal-3*. The plasmids allowing expression of RAD-50 and MRE-11 are from (GIRARD *et al.* 2018).

### Cytology

All analyses were performed on 20-24h post-L4 adults grown at 20°C except for Figure 6, in which the specimen analyzed were 44-48h post L4 adults grown at 15°C.

For immunofluorescence experiments, dissection of gonads, fixation, immuno-staining and DAPI counterstaining were performed as in (MARTINEZ-PEREZ AND VILLENEUVE 2005). The following primary antibodies were used: rabbit anti-HAL-2 (1:10.000, (ZHANG *et al.* 2012)), mouse anti-CYE-1 (1:200, (BRODIGAN *et al.* 2003)), guinea-pig anti-SYP-1 (1:200, (MACQUEEN *et al.* 2002)), rabbit anti-PLK-2 (1:500, (NISHI *et al.* 2008)), rabbit anti-GFP (1:1000, (YOKOO *et al.* 2012)), chicken anti-HTP-3 (1:250, (MACQUEEN *et al.* 2005)), guinea-pig anti-HIM-8 (1:250, (PHILLIPS *et al.* 2005)), rabbit anti-PCH-2 (1:1000, (DESHONG *et al.* 2014)), guinea-pig anti-SUN-1 S8-Pi (1:1000, (PENKNER *et al.* 2009)), rabbit anti-RAD-51 (1:250, (COLAIACOVO *et al.* 2003)), rat anti-PLK-1 (1:50, (LABELLA *et al.* 2011)), rabbit anti-HTP-1/2 (1:500, (MARTINEZ-PEREZ *et al.* 2008)) and rabbit anti-HTP-1 S11-Pi (1:200, (FERRANDIZ *et al.* 2018)). Secondary antibodies were Alexa Fluor 488, 555 and 647-conjugated goat antibodies directed against the appropriate species (1:400, Life Technologies).

For FISH experiments, DNA for 5S rDNA probes was prepared as in (ZHANG *et al.* 2012). Probe DNAs were subsequently labeled with either Alexa Fluor 488 using the Ulysis DNA labeling kit (Life Technologies). Gonad dissection, permeabilization, fixation, hybridization and DAPI counterstaining were performed as described in (NABESHIMA *et al.* 2011).

### Irradiation

Worms were exposed to 5,000 rad (50 Gy) of γ-irradiation using a Cs-137 source at 20 h post-L4 stage. RAD-51 immunostaining was performed on gonads dissected and fixed 1 h after irradiation.

### Image acquisition and processing

High resolution and 3D-Structured Illumination Microscopy images were collected as Z-stacks (at 0.2μm intervals for high resolution and 0.125μm for super-resolution) using a 100x NA 1.40 objective on a DeltaVison OMX Blaze microscopy system, deconvolved and corrected for registration using SoftWoRx. Multiple overlapping fields covering the whole length of the gonad were acquired for each specimen, and gonads were assembled using the “Grid/Collection” plugin (PREIBISCH *et al.* 2009) on Fiji (SCHINDELIN *et al.* 2012). Final assembly of 2D maximum intensity projections was performed using Fiji, with adjustments of brightness and/or contrast made in Fiji or Adobe Photoshop.

### Data and reagent availability

All nematode strains and plasmids generated for this study are available upon request, either from the authors or from the Caenorhabditis Genetics Center.

## Results

### Identification of HAL-3 as a binding partner of HAL-2

To gain further insights into HAL-2 function during meiotic prophase, we pursued a proteomincs approach to identify candidate HAL-2 binding partners using a previously generated transgene expressing a functional GFP::HAL-2 fusion protein in the *hal-2(me79)* null mutant background ((ZHANG *et al.* 2012); Figure S1). We performed immuno-precipitation using an anti-GFP antibody on fractionated nuclear extracts from both wild-type (control) and *hal-2; gfp::hal-2* worms and analyzed the eluates by mass-spectrometry (Figure S1). Under the conditions used, HAL-2 is found almost exclusively in the nuclear soluble fraction (SILVA *et al.* 2014), and accordingly, HAL-2 peptides were detected only in the immunoprecipitate of the nuclear soluble fraction prepared from the strain expressing GFP::HAL-2.

We used available expression data sets and functional annotation to define a list of candidate HAL-2 interactors among proteins with peptides specifically detected in the same fraction as HAL-2 peptides (Figure S1). We then used RNAi to test whether depletion of these candidate gene products resulted in meiotic defects. Indeed, we found that RNAi for *T15H9.2* (hereafter referred to as *hal-3*), led to deficit of crossover-based connections (chiasmata) between homologous chromosomes, detected as an increased number of DAPI-stained bodies in oocytes at diakinesis, the last stage of meiotic prophase (Figure 1A). Further, we used CRISPR mutagenesis to generate *hal-3(me99)*, a 1bp deletion in the first exon of the coding sequence that is predicted to induce a frameshift and an early termination of the translation of the *hal-3* transcript; diakinesis oocytes from the *hal-3(me99)* mutant consistently lack chiasmata (Figure 1A), reflecting underlying defects in early meiotic prophase events that phenocopy loss of *hal-2* function (see below).

**Figure 1:**
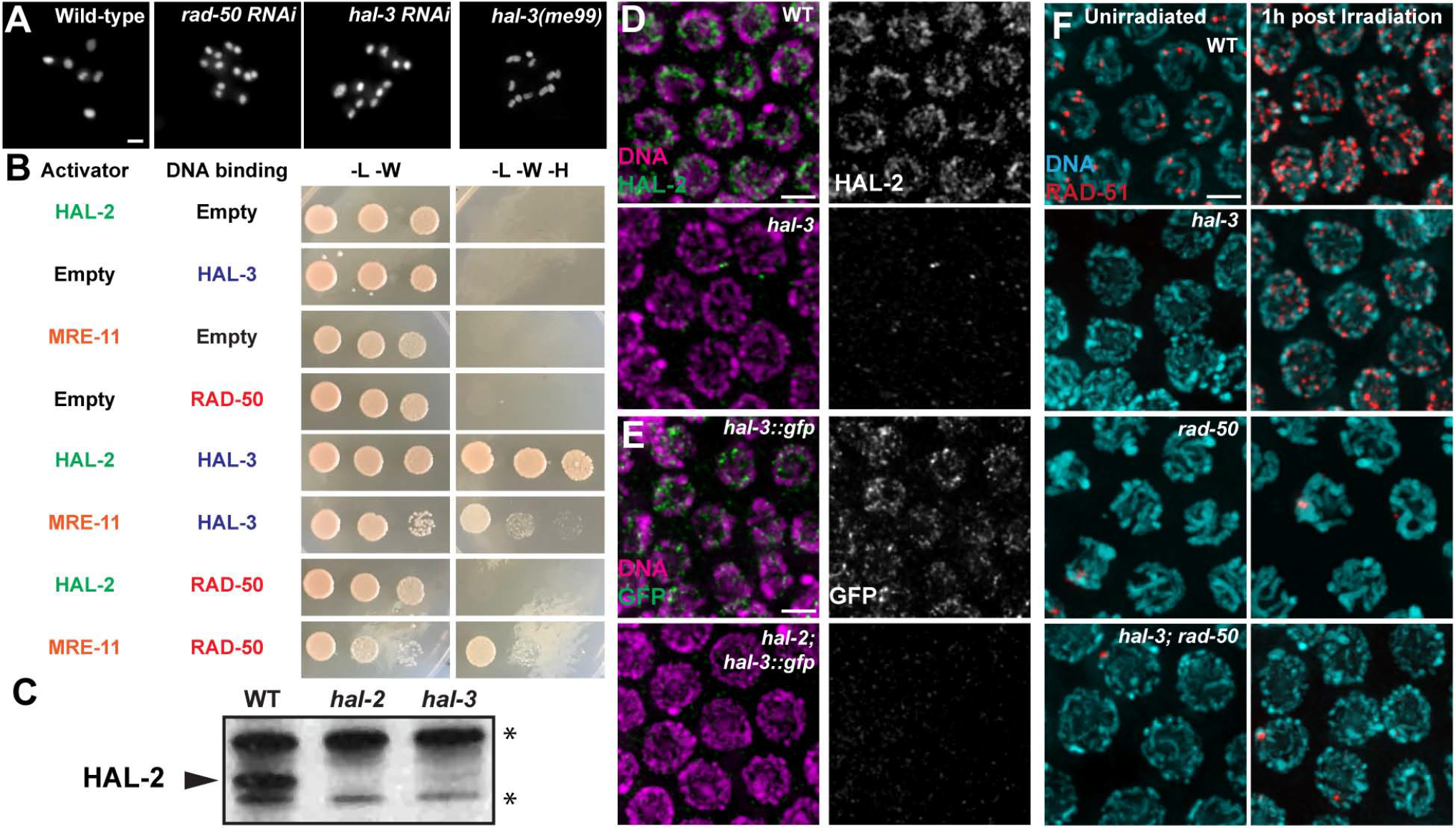
Identification of HAL-3 as an interacting and interdependent partner of HAL-2. **A)**Each panel shows DAPI-stained chromosomes in a single example oocyte nucleus at diakinesis, the last stage of meiotic prophase. Whereas 6 bivalents are present in the wild-type oocyte, each representing a homolog pair held together by a crossover-based connection known as a chiasma, 11 or 12 DAPI bodies are detected in the nuclei from *rad-50 RNAi*, *hal-3 RNAi*, and *hal-3(me99)* worms, reflecting a deficit of chiasmata. **B)** Yeast-two hybrid analysis testing pairwise interaction between HAL-2 and HAL-3 and members of the MRN complex (MRE-11 and RAD-50). Interaction between prey proteins fused with the GAL4 activation domain (Left) and the baits fused with the LexA DNA binding domain (Right) was assayed by growth on media lacking histidine (−LWH). Serial dilutions are spotted (1, 1:100, 1:10,000). **C)** HAL-2 levels, detected by western blotting on whole worm extracts, are severely reduced in *hal-3(me99)* mutant worms. The asterisks indicate non-specific signals detected in extracts of both in wild-type and in *hal-2* and *hal-3* mutant worms **D-E)** Immunodetection of HAL-2 (green, panel D) or HAL-3::GFP (green, panel E) in the nucleoplasm of meiocytes at the mid-pachytene stage of meiotic prophase. HAL-2 and HAL-3::GFP were not detected in *hal-3* and *hal-2* mutants, respectively. **F)** Immunodetection of strand-exchange protein RAD-51 in mid-prophase nuclei from worms of the indicated genotypes, either unirradiated (left) or one hour after exposure to 5kRad of γ-irradiation (right); images illustrate that *hal-3* is required to form meiotic but not radiation-induced RAD-51 foci, and that radiation-induced foci in the *hal-3* mutant are *rad-50* dependent. (A single bright RAD-51 focus is detected in a subset of nuclei in the *rad-50* mutant background, attributable to DNA damage arising during DNA replication (HAYASHI *et al.* 2007). In D-F, images are maximum intensity projections of image stacks encompassing whole nuclei, with DNA counterstained with DAPI. Scale bars in panels A, D, E and F represent 2μm.

A combination of yeast two-hybrid (Y2H), biochemistry and immunolocalization experiments support the conclusion that HAL-2 and HAL-3 are likely subunits of the same protein complex. First, Y2H analysis indicates that HAL-3 interacts with HAL-2 in an orthogonal system (Figure 1B). Second, HAL-2 and HAL-3 are interdependent for localization and/or stability *in vivo* (Figure 1C-E, Figure S2A-B); analysis of whole worm extracts showed that HAL-2 levels were severely reduced in *hal-3(me99)* (Figure 1C) and immunodetection of the HAL-2 protein, which localizes to the nucleoplasm of wild-type germ cells, similarly indicated that HAL-2 was undetectable in the *hal-3* mutant (Figure 1D). Likewise, a HAL-3::GFP fusion protein (expressed from a transgene containing the endogenous *hal-3* regulatory elements) was detected in the nucleoplasm of otherwise wild-type germ cells but was undetectable in the *hal-2* mutant (Figure 1E). Together these data support a scenario in which HAL-2 and HAL-3 function together in a complex that localizes to the nucleoplasm of germ cells and are mutually required for stability and function.

### HAL-2/HAL-3 interacts with MRE-11-RAD-50 complex components but is not required for their functions in DSB repair

Our IP/MS analysis also identified peptides corresponding to RAD-50 specifically in the nuclear soluble fraction containing HAL-2 and HAL-3 peptides (Figure S1C). RAD-50, a component of the MRE-11-RAD-50 complex, was previously shown to be required both for initiation of meiotic recombination through the formation of double-strand DNA breaks (DSBs) and for resection of DSBs to yield 3’ ssDNA tails that can engage in homologous recombination-based repair (HAYASHI *et al.* 2007). We did not detect a direct interaction between HAL-2 and RAD-50 by Y2H assay, but we did detect an interaction between HAL-3 and MRE-11 and confirmed previously-reported Y2H interactions between MRE-11 and RAD-50 (YIN AND SMOLIKOVE 2013; GIRARD *et al.* 2018), suggesting that the HAL-2/HAL-3 and MRE-11-RAD-50 complexes may indeed interact *in vivo*.

Despite this evidence for physical interactions between HAL-2/HAL-3 and MRE-11-RAD-50 complexes, cytological analysis of RAD-51 foci, which mark the sites of processed DSBs, revealed that HAL-3 is not required for function of MRE-11-RAD-50 in DSB resection during meiotic prophase (Figure 1F, Figure S2C). As described previously for *hal-2* mutants (ZHANG *et al.* 2012), *hal-3* mutants exhibited a strong reduction in meiotic RAD-51 foci in early pachytene, where they are abundant in wild-type (Figure 1F, left). However, RAD-51 foci were detected throughout the germline of *hal-3* mutant worms 1 h following exposure to ionizing radiation (IR) (Figure 1D, right; Figure S2C), indicating that DNA damage can be processed efficiently during early meiotic prophase in the absence of HAL-3. Together, these findings suggest that HAL-3 (like HAL-2) is required for the formation of SPO-11-induced DSBs during meiosis but is not required for DSB resection if DSBs are formed by other means. This contrasts with RAD-50, which is required both for SPO-11-dependent DSB formation and for resection of IR-induced DSBs during early meiotic prophase (HAYASHI *et al.* 2007). Moreover, formation of IR-induced RAD-51 foci in early prophase in the *hal-3* mutant is dependent on RAD-50, as such foci were not detected in early prophase in the *hal-3; rad-50* double mutant. Thus, while both HAL-3 and RAD-50 are required for meiotic DSB formation, we infer that the ability to form IR-induced RAD-51 foci in early prophase in the *hal-3* single mutant reflects presence of the resection-promoting activity of MRE-11-RAD-50.

### Multiple aspects of meiotic chromosome organization are impaired in *hal-3* mutants

*hal-3* mutants exhibit severe defects in multiple aspects of meiotic nuclear organization, comparable to those previously reported for mutants lacking HAL-2 ((ZHANG *et al.* 2012); Figure 2). First, DAPI staining indicated that germ cells entering meiotic prophase lack the dramatic reorganization of chromosomes into a clustered configuration characteristic of wild-type meiosis (Figure 2A). This lack of nuclear reorganization is not likely to be due to defective transition from mitosis to meiosis, as we observed that the disappearance of Cyclin E1 (CYE-1, a marker of mitotically cycling cells) and the coalescence of meiotic chromosome axis component HTP-3 into thread-like structures (indicative of entry into meiotic prophase) occurred at comparable positions within the gonads of wild type and *hal-3* and *hal-2* mutants (Figures 2A and S3). However, impairment of nuclear reorganization is correlated with a failure to establish pairing between homologous chromosomes in the *hal-3* mutant, as indicated by failure to detect pairing at both the 5S rDNA locus on chromosome V (Figure 2B) and the X chromosome Pairing Center (PC) at the left end of the X chromosome (Figure 2C).

**Figure 2:**
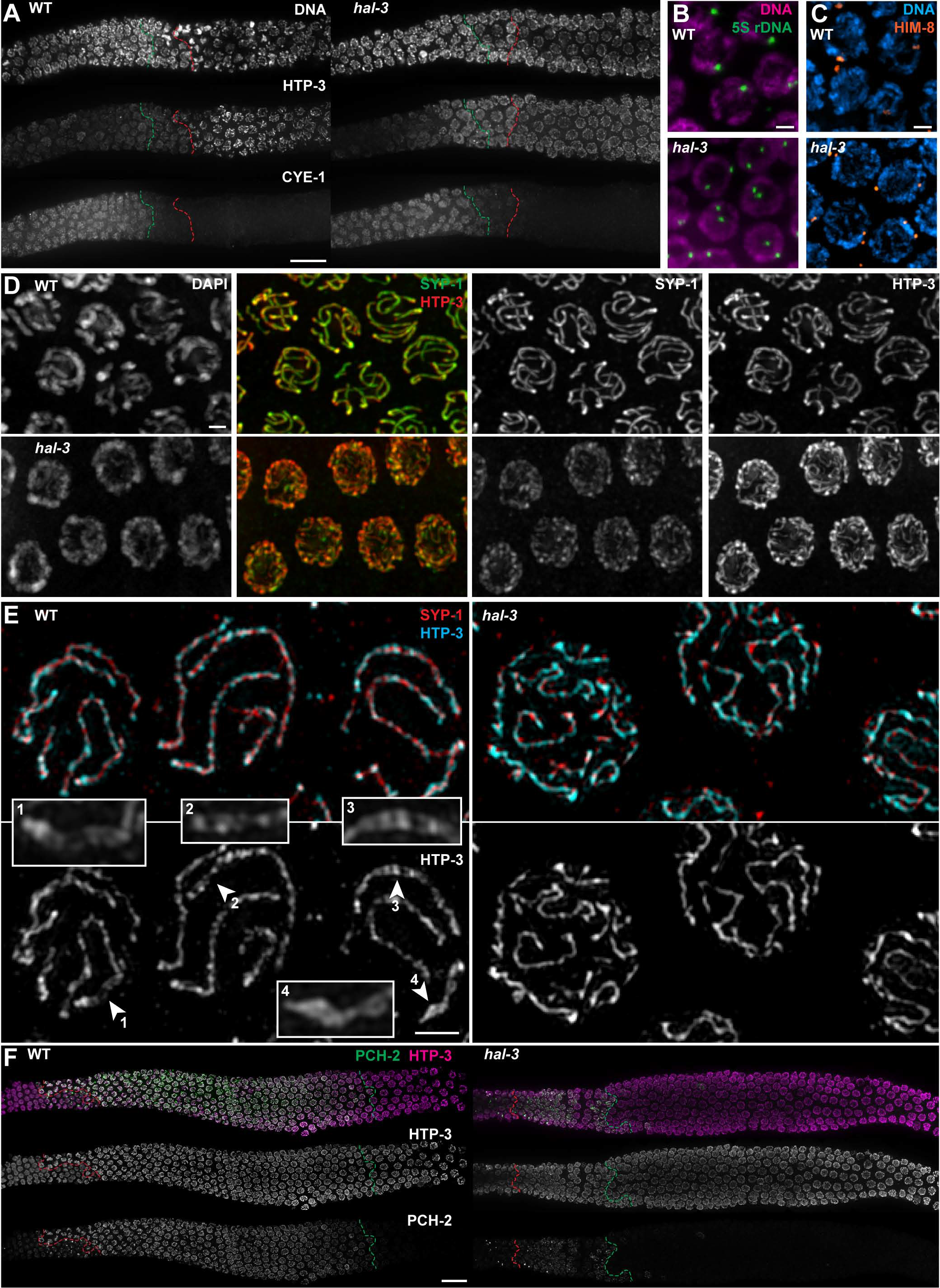
Impaired meiotic prophase chromosome organization in the *hal-3* mutant. **A)**Images of portions of gonads spanning the region where nuclei are exiting the mitotic cell cycle, as indicated by loss of Cyclin E1 (CYE-1) (green dashed line) and entering meiotic prophase, as indicated by coalescence of meiotic chromosome axis component HTP-3 into discrete axial tracks (red dashed line). The relative positions of loss of CYE-1 and acquisition of axial HTP-3 organization are comparable in WT and the *hal-3* mutant; however, the clustering and asymmetric organization of DAPI-stained DNA, reflecting chromosome movements associated with meiotic prophase entry in wild type, is absent in the *hal-3* mutant. **B&C)** Assessment of pairing at the 5S rDNA locus using FISH (green, panel B) and at the X chromosome PC using immunodetection of the X chromosome PC binding protein HIM-8 (orange, panel C) reveals two separate 5S FISH or HIM-8 signals in mid-prophase nuclei in the *hal-3(me99)* mutant (in contrast to single signals present in wild-type nuclei), reflecting failure of homolog pairing in the mutant. **D)** High-resolution deconvolved wide-field images of meiotic chromosome axis component HTP-3 (red) and SC-CR component SYP-1 (green) in fields of nuclei from the mid-pachytene regions of gonads of the indicated genotypes. HTP-3 and SYP-1 colocalize at the interface between paired homologs in wild type meiocytes, whereas in *hal-3* meiocytes, SYP-1 localizes along HTP-3 tracks corresponding to unpaired chromosomes. **E)** 3D-SIM images of nuclei from the same samples as in panel D; images are partial projections through the upper half of the depicted nuclei. Under these imaging conditions, the tripartite organization of the SC can be detected in some segments as two resolvable HTP-3 tracks flanking a SYP-1 signal (arrowheads and corresponding insets). Such tripartite organization was not observed in the *hal-3* mutant nuclei, indicating that SYP-1 is associated with individual chromosome axes. **F)** Detection of the synapsis checkpoint protein PCH-2 (green) and axis component HTP-3 (magenta) in gonads of the indicated genotypes, illustrating premature loss of PCH-2 in the *hal-3* mutant. Red dashed lines indicate the position of the onset of meiotic prophase (identified by the presence of assembled HTP-3 marked chromosomes axes); green dashed lines indicate the end of the zone of nuclei with detectable PCH-2 levels. In panels A, B, C, D and F, images are maximum intensity projections of image stacks encompassing nuclei in the top half of the gonad. Scale bar represents 20μm in panels A and F, and 2μm in panels B, C, D and E.

During wild-type meiosis, pairing is stabilized by the SC, a structure in which components of the SC central region assemble exclusively between aligned homologs, linking the axes of the homologs in parallel along their entire lengths. In contrast, SC-CR proteins were shown to load anomalously in *hal-2* mutants, inappropriately associating with individual, unpaired chromosome axes (ZHANG *et al.* 2012). We therefore analyzed the distribution of SC-CR component SYP-1 and axis component HTP-3 in whole mount gonads of *hal-3* mutants. Analysis of high-resolution images revealed a phenotype identical to that described for *hal-2* mutants: SYP-1 was detected in association with HTP-3 tracks, but the number and relative intensity of HTP-3 signals suggested that these structures contained a single unpaired axis (Figure 2D). This was confirmed using 3D-structured illumination microscopy (SIM) (Figure 2E). In SIM images of wild-type pachytene nuclei, the two separate axes of the aligned homologs are often detected as two resolvable HTP-3 tracks flanking a SYP-1 signal (Figure 2E, arrowheads), whereas in SIM images of *hal-3* mutant nuclei, SYP-1 signals are consistently observed along individual HTP-3 tracks.

In addition to assessing the status of homolog pairing and SC assembly in *hal-3* mutants, we also analyzed the localization of PCH-2, an AAA+ ATPase that has been proposed to play roles in quality control of synapsis (BHALLA AND DERNBURG 2005; DESHONG *et al.* 2014). During wild-type meiosis, PCH-2 associates with SCs in a *syp*-dependent manner at the onset of the pachytene stage and remains associated with SCs until after nuclei have transitioned to late pachytene ((DESHONG *et al.* 2014) and Figure 2F). In *hal-3* mutants, PCH-2 similarly associates with chromosomes soon after SYP loading, but dissociates from chromosomes prematurely (Figure 2F).

### Mis-localization of Polo-like kinases in *hal* mutants

The Polo-like kinases have recently emerged as important regulators of multiple aspects of the meiotic program, including pairing, synapsis, recombination and protection of sister-chromatid cohesion (HARPER *et al.* 2011; LABELLA *et al.* 2011; MACHOVINA *et al.* 2016; NADARAJAN *et al.* 2017; PATTABIRAMAN *et al.* 2017; FERRANDIZ *et al.* 2018). We therefore examined how their localization was affected by loss of *hal-3* function.

We first revisited the dynamic localization of the major meiotic Polo-like kinase PLK-2 in wild-type germ lines (Figure 3A). As previously reported (HARPER *et al.* 2011; LABELLA *et al.* 2011), PLK-2 became concentrated in bright foci and patches at the nuclear envelope (NE) upon entry into meiotic prophase; these NE-associated foci and patches illustrate the localization of PLK-2 at chromosome-NE attachment sites, where PLK-2 promotes PC-led chromosome movements that are important for homolog pairing and synapsis. NE localization of PLK-2 was lost by the end of early pachytene, and PLK-2 subsequently became localized to the SCs. In addition to these previously reported localization patterns, however, we also detected a strong nucleoplasmic localization of PLK-2 in premeiotic germ cell nuclei (Figure 3A). This implies that entry into meiotic prophase triggers a major reorganization of the localization of PLK-2 from the nucleoplasm to chromosome-NE attachment sites.

**Figure 3:**
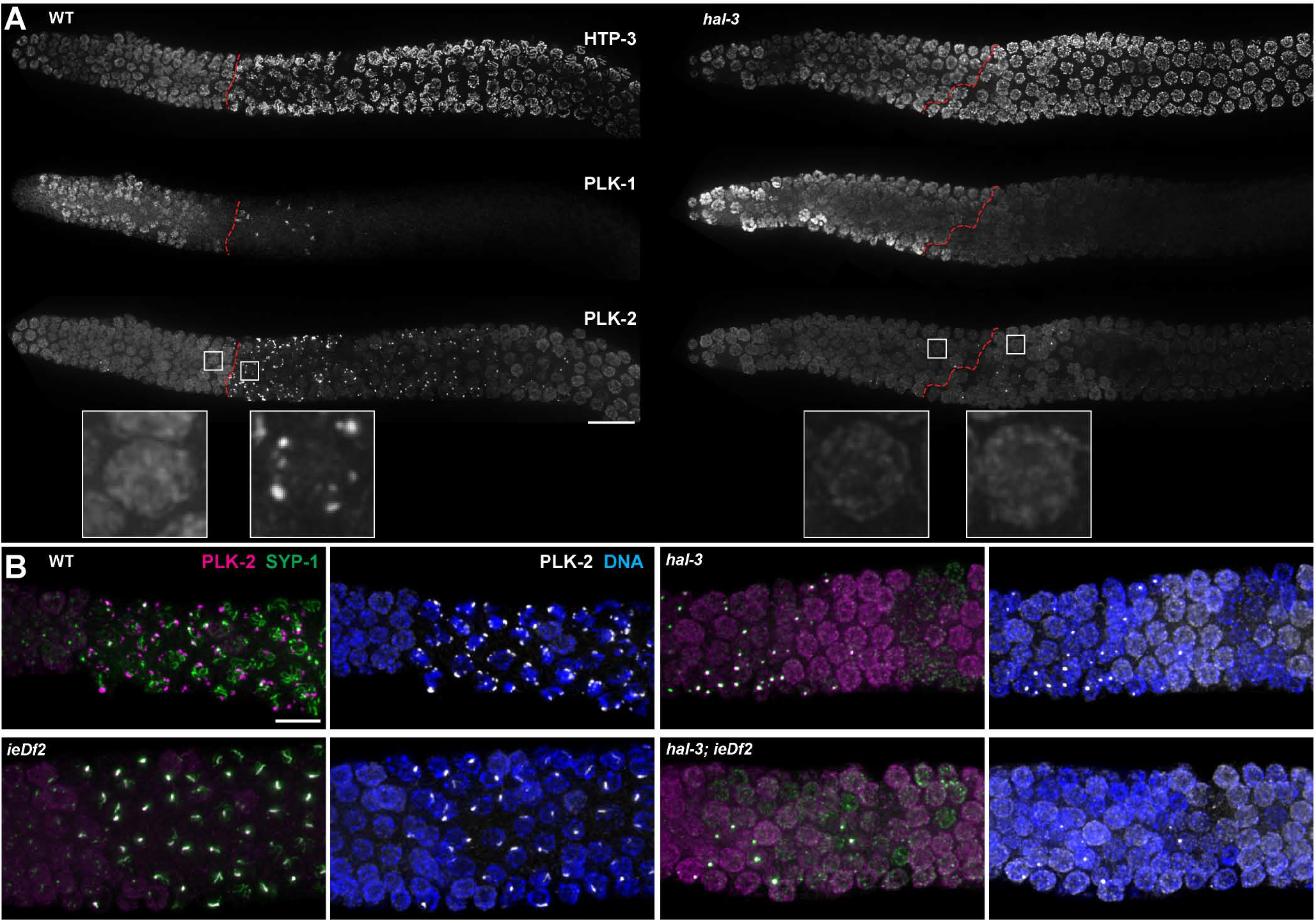
Polo-like kinases are mis-localized in hal mutants. **A)** Immunodetection of Polo-like kinases PLK-1 and PLK-2 in the distal region of the germ line; the red dashed line indicates the onset of meiotic prophase, as identified based on the presence of assembled HTP-3-bound chromosome axes. Insets provide higher magnification of single nuclei located on either side on meiotic prophase entry. In the wild-type gonad, the PLK-1 signal diminishes and is lost from most nuclei prior to meiotic entry, and the PLK-2 signal quickly relocalizes from the nucleoplasm to nuclear-envelope associated foci upon entry into prophase. In the *hal-3* mutant gonad, PLK-2 does not relocalize to NE-associated foci but instead remains strongly nucleoplasmic following meiotic prophase entry, and PLK-1 can still be detected in the early prophase nuclei. Scale bars represent 20μm. **B)** Comparison of effects of loss of *hal-3* function and removal of all PC-binding proteins (caused by the deletion *ieDf2*) on PLK-2 localization in the region of the gonad where nuclei enter meiotic prophase. The left column illustrates localization of PLK-2 (purple) relative to SYP-1 (green) and the middle column depicts localization of PLK-2 (white) relative to DAPI-stained chromatin (blue). In wild-type, PLK-2 localizes to NE-associated chromosome attachment sites, from which stretches of SYP-1 emerge, reflecting SC assembly between paired homologs, initiated at PCs. In the *ieDf2* germ line, DAPI-stained chromatin remains broadly distributed around the nuclei following meiotic entry (similar to the *hal-3* mutant), reflecting absence of PC-binding proteins and consequent lack of PC-mediated chromosome movement. However, in contrast to persistence of PLK-2 as a diffuse nucleoplasmic signal as in the *hal-3* mutant (and *hal-3; ieDf2* double mutant), PLK-2 is instead concentrated to a single bright signal at the nuclear periphery that colocalizes with an aggregate of SYP-1. Scale bars represent 10μm.

We also examined the localization of PLK-1, another Polo-like kinase expressed in the *C. elegans* germ line (Figure 3A). PLK-1 is primarily required for meiotic spindle function, nuclear envelope breakdown in mitosis, and embryonic polarity (CHASE *et al.* 2000; NISHI *et al.* 2008; RIVERS *et al.* 2008), but it has also been shown to be able to partially compensate for loss of PLK-2 function during meiotic prophase (HARPER *et al.* 2011; LABELLA *et al.* 2011). In wild-type gonads, PLK-1 signal was detected in premeiotic germ cell nuclei, exhibiting chromosome adjacent and/or nucleoplasmic localization, and it became largely undetectable (except for an occasional patch at the nuclear periphery) as nuclei transitioned into meiotic prophase.

*hal-3* mutants exhibited several anomalies in the localization of these Polo-like kinases (Figure 3A). As previously reported for *hal-2* (ZHANG *et al.* 2012), PLK-2 failed to localize to discrete foci and patches upon meiotic entry. Moreover, strong nucleoplasmic PLK-2 signals were instead detected for multiple cell rows beyond meiotic entry (Figure 3A). PLK-1 likewise exhibited anomalous localization in *hal-3* mutant germ lines, as detectable levels of nucleoplasmic and/or chromosome adjacent PLK-1 were observed in the nuclei of some germ cells that had entered meiotic prophase (based on the presence of coalesced chromosome axes, Figure 3A). Altogether, these data indicate that *hal-3* mutant meiocytes have abnormally high levels of Polo-like kinases in their nucleoplasm following meiotic entry.

### Absence of early NE-associated PLK phosphorylation targets is not sufficient to relocalize PLK-2 to the nucleoplasm

As described above, PLK-2 associates with chromosome-NE attachment sites upon entry into meiotic prophase in wild-type germ cells, and previous work has demonstrated that this association is a consequence of the binding of PLK-2 to a family of zinc-finger proteins, HIM-8 and ZIM-1,−2 and −3, that localize with PCs (PHILLIPS *et al.* 2005; PHILLIPS AND DERNBURG 2006; HARPER *et al.* 2011; LABELLA *et al.* 2011; KIM *et al.* 2015). We therefore tested whether the distribution of PLK-2 observed in *hal* mutants could be mimicked by absence of these PC-binding proteins (Figure 3B). Specifically, we analyzed the distribution of PLK-2 in worms homozygous for a deficiency (*ieDf2*) engineered to delete the entire cluster of genes encoding these proteins (HARPER *et al.* 2011). Despite absence of these primary early meiotic targets, PLK-2 did not become dispersed in the nucleoplasm in *ieDf2* homozygotes but instead became localized to aggregates containing SC-CR protein SYP-1 (polycomplexes) that form in this context (Figure 3B). This indicates that the nucleoplasmic localization of PLK-2 observed in early prophase nuclei in *hal-3* and *hal-2* mutants is not simply a consequence of lack of interaction with its normal early prophase targets and instead indicates a more active role for HAL-3 in antagonizing the nucleoplasmic localization of PLK-2.

In *ieDf2* homozygotes, SYP-1 eventually becomes colocalized with a subset of chromosome axis segments, reflecting foldback synapsis linking the axes of non-homologously “paired” segments of chromosomes that had failed to associate with their homologs ((HARPER *et al.* 2011); Figure S4). However, in *hal-3; ieDf2* double mutants, SYP-1 is instead detected all along the lengths of individual unpaired axes, as in hal single mutants. Thus, the aberrant localization behavior of SYP-1 observed when PC binding proteins are absent is nevertheless governed by the HAL-2/HAL-3 complex.

### Reduction of CHK-2 activity contributes to the mis-localization of PLK-2 in *hal-2/3* mutants

Association of PLK-2 with PCs upon entry into meiotic prophase is dependent on CHK-2-mediated phosphorylation of PC-binding proteins (HARPER *et al.* 2011). As CHK-2 activity was previously reported to be defective in the *hal-2* mutant (ZHANG *et al.* 2012), we wondered whether the anomalous localization of PLKs in the nucleoplasm of *hal-2/3* mutant meiocytes could be due to reduced activity of CHK-2. Thus, we analyzed the localization of PLK-2 in *chk-2* mutant worms (Figure 4A). We observed that in *chk-2* mutant germ cells, similarly to what we had observed in *hal* mutants, PLK-2 not only fails to concentrate at chromosome-NE attachment sites upon meiotic entry but also remains diffuse in the nucleoplasm. This suggests that the mis-localization of PLK-2 in *hal* mutants could be due to defective CHK-2 activity.

**Figure 4:**
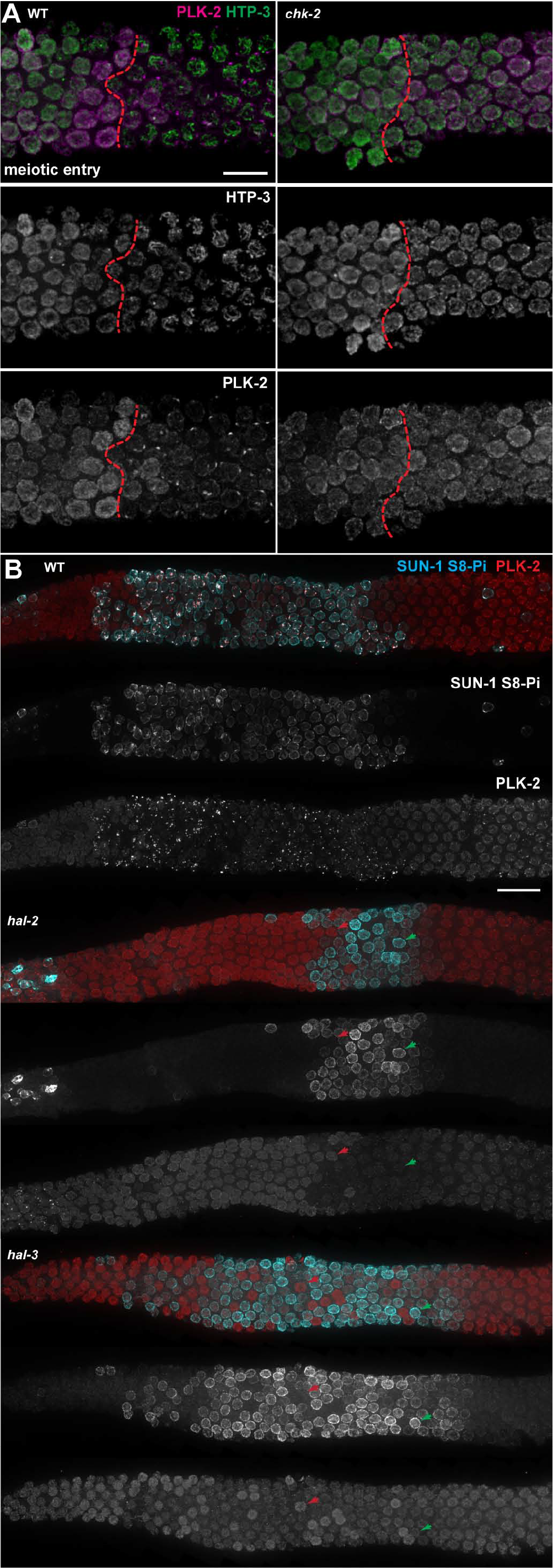
Impact of CHK-2 activity on the localization of PLK-2. **A)** Immunodetection of axis component HTP-3 (red) and Polo-like kinase PLK-2 in the region of the gonad encompassing entry into meiotic prophase of the indicated genotypes, showing that (as in *hal* mutants) levels of nucleoplasmic PLK-2 remain high following meiotic prophase entry in *chk-2* mutants. Dashed lines indicate entry into meiotic prophase based on coalescence of chromosome axes. **B)** Simultaneous immunostaining for PLK-2 and NE protein SUN-1 S8-Pi (a marker of CHK-2 activity) in portions of gonads spanning from just prior to meiotic entry (left) through mid-prophase, illustrating the impact of CHK-2 activity on the localization of PLK-2 in *hal* mutants. In the wild-type, the presence of SUN-1 S8-Pi in meiotic prophase nuclei is strongly correlated with the localization of PLK-2 at NE foci, whereas in the *hal* mutant gonads, levels of nucleoplasmic PLK-2 signals anti-correlate with SUN-1 S8-Pi signals; extreme examples are indicated by red and green arrowheads. For both A and B, images are maximum intensity projections of image stacks encompassing nuclei in top half of the gonad; scale bars represent 20μm.

We evaluated the status of CHK-2 activity by assessing phosphorylation on Serine 8 of the NE protein SUN-1 (SUN-1 S8-Pi), which serves as a marker of early prophase CHK-2 activity (PENKNER *et al.* 2009). This analysis showed that CHK-2 activity is indeed reduced in most *hal-3* mutant gonads (Figure S5). However, as in the *hal-2* mutant, variable numbers of nuclei with residual SUN-1 S8-Pi staining were observed in *hal-3* mutant gonads ((ZHANG *et al.* 2012) and Figures 4B and S5), often interspersed with nuclei lacking detectable SUN-1 S8-Pi. This created a unique opportunity to examine the relationship between PLK-2 localization and CHK-2 activity within different nuclei from the same gonad, by comparing PLK-2 immunostaining (in *hal-2* and *hal-3* mutants) in nuclei with and without detectable SUN-1 S8-Pi (Figure 4B). We observed that, in our preparations, nuclei with low or undetectable SUN-1 S8-Pi signals tended to have higher levels of nucleoplasmic PLK-2, and conversely, that higher levels of SUN-1 S8Pi were detected in nuclei with low levels of PLK-2. This implies that, in *hal* mutants, residual CHK-2 activity either triggers the degradation of PLK-2 or promotes its recruitment to some unknown target that is washed out during staining procedure. We note, however, that the residual CHK-2 activity detected in *hal* mutants is insufficient to promote association of PLK-2 to its wild-type targets (the PCs or the SCs), and that even in nuclei with detectable CHK-2 activity, a pool of PLK-2 was consistently detected in the nucleoplasm of *hal* mutant meiocytes.

### Abnormal PLK activity in *hal* mutants

Our observations that PLKs are mis-localized and that elevated nucleoplasmic PLK protein is correlated with defective acquisition of early-pachytene features in *hal* mutants led us to hypothesize that some of the defects might be due to inappropriate/untimely activity of PLKs on substrates from which they would normally be sequestered. To test this hypothesis, we analyzed the impact of loss of *hal-2* or *hal-3* on the phosphorylation of Serine 11 of the chromosome axis components HTP-1 (HTP-1 S11-Pi), which was recently shown to be phosphorylated during meiotic prophase in a PLK-1/2 dependent manner (FERRANDIZ *et al.* 2018). HTP-1 is found both in the nucleoplasm and bound to chromosome axes during early meiotic prophase (SILVA *et al.* 2014) and thus might potentially be affected by the mis-localization of PLKs observed in *hal* mutants.

As HTP-1 loading on chromosome axes was previously reported to be partially defective in *hal-2* (ZHANG *et al.* 2012), analysis of HTP-1 phosphorylation in the *hal-2* mutant background was carried out in a strain in which the sole source of HTP-1 was a transgene expressing a GFP fusion protein, allowing simultaneous detection of both total HTP-1 (using an anti-GFP antibody) and the fraction that is phosphorylated (using an anti-HTP-1 S11-Pi antibody; (FERRANDIZ *et al.* 2018)). HTP-1 S11-Pi immunofluorescence signals on chromosome axes were comparable in *hal-2* mutants and controls even though overall axis-associated HTP-1 signals were greatly diminished in the *hal-2* mutant. Thus, the ratio of phosphorylated to total HTP-1 on chromosome axes (defined by the presence of the HTP-3 protein) in early pachytene meiocytes was reliably higher in *hal-2* mutants than in control germ lines (Figure 5A). We likewise observed comparable HTP-1 S11-Pi signals in wild-type and *hal-3* mutant early pachytene meiocytes (Figure 5B) despite an overall reduction in axis-associated HTP-1 in the *hal-3* mutant (Figure S6). Together, these data indicate that a nuclear PLK-1/2 target is hyper-phosphorylated on chromosome axes in *hal* mutants.

**Figure 5:**
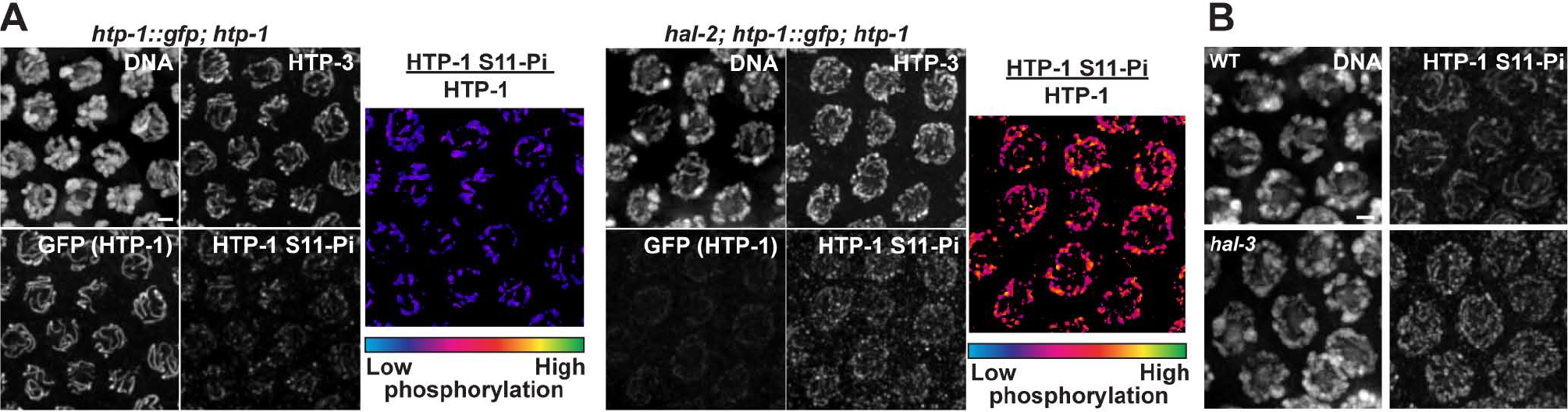
Inappropriate phosphorylation of Polo-like kinase substrate HTP-1 S11 in *hal* mutants. **A)** Grayscale panels: Simultaneous immunodetection of HTP-3, HTP-1::GFP, and HTP-1 S11-Pi in control and *hal-2* mid-pachytene nuclei in a genetic background in which the sole source of HTP-1 is a transgene expressing HTP-1::GFP. Images show comparable HTP-1 S11-Pi signals in mutant and control despite very low levels of total axis-associated HTP-1 (detected using anti-GFP antibody) in the mutant. Color panels: representation of the relative phosphorylation levels at S11 of axis-associated HTP-1, using the “Fire” lookup table to illustrate the elevated ratio of HTP-1 S11-Pi / total HTP-1 associated with meiotic chromosome axes in the *hal-2* mutant relative to control. For this display, the ratio of anti-HTP-1 S11-Pi / anti-GFP signals was calculated and represented for each position identified as chromosome axis based on the presence of HTP-3, using the same scale for both genotypes. B) Detection of comparable HTP-1 S11-Pi immunofluorescence signals on chromosome axes in wild-type and *hal-3* mutant mid-pachytene nuclei, despite detection of severely reduced axis-associated HTP-1 in the *hal-3* mutant in separate experiments (Figure S6). Images in A and B are maximum intensity projections of image stacks encompassing nuclei in the top half of the gonad; scale bars represent 2μm.

### Loss of *plk-2* function partially rescues meiotic defects in *hal-2* mutants

Our observations of mis-localized PLKs and evidence for untimely phosphorylation of a PLK substrate in *hal* mutants suggested that at least some of the defects in meiotic chromosome organization could be due to inappropriate PLK activity. We further tested this hypothesis by analyzing the impact of the deletion of *plk-2* on the *hal-2* mutant phenotype. We first examined the localization of axis component HTP-3 and SC-CR protein SYP-1 in the *hal-2; plk-2* double mutant, which revealed that loss of *plk-2* function partially restores the ability to assemble tripartite SCs in the *hal-2* mutant background (Figure 6A). Although homolog pairing and synapsis are partially impaired in *plk-2* null mutants, by mid-prophase *plk-2* mutant nuclei harbor a mixture of synapsed and unsynapsed chromosome axes (HARPER *et al.* 2011; LABELLA *et al.* 2011) and SIM imaging revealed normal-appearing tripartite SCs (Figure 6A). As in the *hal-3* single mutants, tripartite SC structure was not detected in the *hal-2* single mutant, and the SYP-1 signal typically had a punctate, discontinuous appearance along the lengths of individual unpaired axes (Figure 6A, cyan boxes). In contrast, nearly all mid-pachytene nuclei in *hal-2; plk-2* gonads contained at least one stretch of continuous SYP-1 staining, and SIM imaging revealed that these continuous stretches of SYP-1 reflect the presence of tripartite SCs in which SC-CR components were localized between a pair of resolvable HTP-3 tracks corresponding to two separate chromosome axis segments (Figure 6A, magenta boxes). As loss of *plk-2* function restored segments of tripartite SC in the *hal-2* mutant background, we infer that PLK-2 activity is at least partially responsible for the inability of SYP proteins to link chromosome axes in *hal* mutants. However, the *hal-2; plk-2* double mutant also displayed a feature typical of the *hal-2/3* single mutants but not observed in the *plk-2* single mutant, *i.e.* association of SYP-1 along unpaired chromosome axes (Figure 6A, cyan boxes). Thus, absence of PLK-2 alone is not sufficient to fully rescue the SC assembly defects caused by loss of HAL-2/HAL-3.

**Figure 6:**
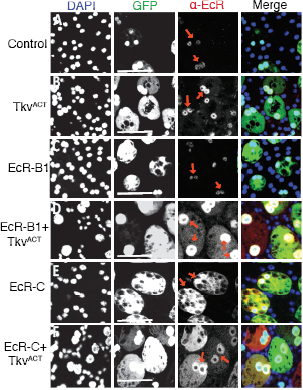
A mutation in *plk-2* partially rescues *hal-2* meiotic defects. **A)** 3D-SIM images of axis component HTP-3 and SC-CR protein SYP-1 in nuclei from the mid-prophase region of gonads of the indicated genotypes; images are partial projections through the upper half of the depicted nuclei. Magenta arrowheads and corresponding enlarged insets highlight examples of SC segments where tripartite SC organization is detected as pairs of spatially-resolved HTP-3 tracks with SYP-1 signal located between them. Cyan arrowheads and corresponding enlarged insets highlight examples where SYP-1 is detected on an unpaired axis segment. White arrowhead indicates an example of a chromosome axis lacking SYP-1 in the *plk-2* single mutant. In contrast to the *hal-2* single mutant, tripartite SC stretches are detected in the *hal-2; plk-2* double mutant, **B)** Detection of the synapsis-checkpoint protein PCH-2 (green) and axis component HTP-3 (magenta) in gonads of the indicated genotypes. Red dashed lines indicate the position of the onset of meiotic prophase; green dashed lines indicate the end of the zone of nuclei with detectable PCH-2 levels. Scale bar represents 20μm.

We also examined how the distribution of the PCH-2 protein was affected by loss of *plk-2* function (Figure 6B). We found that PCH-2: *i*) remained associated with chromosomes throughout the entire pachytene zone in the *plk-2* single mutant, *ii*) was lost prematurely in the *hal-2* single (as in the *hal-3* single mutant), and *iii*) exhibited an intermediate phenotype in the *hal-2; plk-2* double mutant. These observations together indicate that PLK activity promotes premature loss of PCH-2 when *hal*-2/3 function is compromised, and conversely, that the HAL-2/HAL-3 complex promotes prolonged retention of PCH-2 when *plk-2* function is compromised.

## Discussion

In this study, we identified HAL-3 as a partner of HAL-2, a factor previously implicated in the coordination of early prophase events during *C. elegans* meiosis. Analysis of the *hal-3* mutant phenotype revealed that localization of HAL-2 in meiocytes is dependent on HAL-3, and accordingly, that the meiotic defects of the *hal-3* mutant are virtually indistinguishable from those previously described for the *hal-2* mutant: defective homolog pairing and loading of SC-CR components onto unpaired chromosome axes. Conversely, cytological analysis of a strain expressing a HAL-3::GFP fusion protein indicated that HAL-2 is required for the localization of HAL-3. These observations and additional yeast two-hybrid experiments are consistent with a model in which HAL-2 and HAL-3 form a complex *in vivo* that localizes to the nucleoplasm of meiotic nuclei and participates in the coordination of prophase events.

Our analysis of the *hal-3* mutant highlights a previously unappreciated consequence of the loss of *hal-2/3* function on the biology of Polo-like kinase during meiotic prophase. We confirm and extend the observation that concentration of PLK-2 at NE-associated PCs during early meiotic prophase is dependent on the function of HAL-2/HAL-3 (ZHANG *et al.* 2012). Importantly, we show that PLK-2 not only fails to be localized correctly in *hal-2/3* mutants but is in fact mis-localized: PLK-2 (and PLK-1) remain at higher than normal levels in the nucleoplasm of early prophase meiocytes in *hal-2/3* mutants. Furthermore, deletion of the major meiotic Polo-like kinase, PLK-2, is sufficient to rescue some of the meiotic defects observed in *hal-2/3* mutants, thereby supporting a model in which the HAL-2/HAL-3 complex regulates early meiotic prophase events, at least in part, by effectively preventing association of PLKs with untimely targets.

### Role of HAL-2 -HAL-3 in regulation of meiotic prophase events by Polo-like kinases

Following the discovery of the requirement for Polo in the correct assembly of mitotic spindles in Drosophila (SUNKEL AND GLOVER 1988), Polo-like kinases have been recognized as master regulators of many aspects of cell cycle progression, including centriole biology, cytokinesis and DNA damage response (reviewed in (ZITOUNI *et al.* 2014)). Roles of Polo-like kinases as key regulators of major cell-cycle transitions have also been observed in meiosis, as illustrated in *S. cerevisiae*, in which exit from pachytene is driven by expression of the Polo-like kinase CDC5p (SOURIRAJAN AND LICHTEN 2008) or in *D. melanogaster* oogenesis, where maintenance of the lengthy G2 arrest that follows pachytene and precedes nuclear envelope breakdown is controlled by Matrimony, an inhibitor of Polo (XIANG *et al.* 2007).

In the context of *C. elegans* meiosis, Polo-like kinases have emerged as major regulators of multiple distinct meiotic prophase events rather than as regulators of cell-cycle transitions *per se*. Polo-like kinases govern many essential processes of meiotic prophase including homolog pairing, synapsis, recombination and chromosome segregation (HARPER *et al.* 2011; LABELLA *et al.* 2011; MACHOVINA *et al.* 2016; NADARAJAN *et al.* 2017; PATTABIRAMAN *et al.* 2017; FERRANDIZ *et al.* 2018). This regulatory action of Polo-like kinases on so many distinct processes is thought to be governed by their dynamic localization during prophase progression, as PLK-2 (the major meiotic PLK) is initially found upon meiotic prophase entry at the chromosome-NE attachments sites, recruited by the PC binding proteins to promote pairing and synapsis (HARPER *et al.* 2011; LABELLA *et al.* 2011), then relocalizes in a recombination-dependent manner during mid-pachytene to SCs, where it alters the SC dynamics and negatively regulates initiation of meiotic recombination (MACHOVINA *et al.* 2016; NADARAJAN *et al.* 2017; PATTABIRAMAN *et al.* 2017). Finally, PLK-2 becomes concentrated on a subdomain (known as the short arm) of each bivalent during late meiotic prophase to enable the two-step release of sister chromatid cohesion across the two meiotic divisions (FERRANDIZ *et al.* 2018; SATO-CARLTON *et al.* 2018).

How is this dynamic association to nuclear structures regulated during prophase? Studies on the role of Polo-like kinases during the mitotic cell cycle indicate that their localization is regulated at least in part through interactions of a domain of PLK called the Polo Box Domain (PBD) with proteins that contain a consensus binding motif Ser-[pSer/pThr]-[Pro/X], phosphorylated by the action of a priming kinase (reviewed in (ZITOUNI *et al.* 2014)). Two recent studies have demonstrated that Polo-like kinase localization during meiotic prophase in *C. elegans* similarly relies on such interactions with phosphorylated targets. Concentration of PLK-2 to the short arm of the bivalent during late prophase is dependent on phosphorylation of a Polo-binding motif in the C-terminus of SC-CR component SYP-1 (SATO-CARLTON *et al.* 2018). Moreover, upon meiotic entry, PLK-2 binds to conserved polo-binding motifs on the PC-binding proteins, phosphorylated by the priming kinase CHK-2 (KIM *et al.* 2015).

The current work shows that HAL-2/HAL-3 also governs the localization of PLK-2 upon meiotic entry. It is unlikely that HAL-2/HAL-3 regulates PLK localization primarily by modifying the availability of PC-binding proteins, as deletion of the operon encoding all four of the PC-binding proteins does not result in high nucleoplasmic PLK-2 and leads to meiotic defects that are distinct from those observed in *hal-2/3* mutants ((HARPER *et al.* 2011) and this study). Instead, retention of high levels of PLK-2 in the nucleoplasm in *hal* mutants appears to result, at least in part, from reduced activity of CHK-2. However, a high level of nucleoplasmic PLK-2 (resulting from reduced CHK-2 activity) is not sufficient to explain the mis-regulation of PLK function observed in *hal* mutants, as the defects in meiotic chromosome organization in *hal-2/3* mutants are distinct from those observed in *chk-2* null mutants. We therefore propose the HAL-2/HAL-3 complex must play an additional role in regulating PLK activity beyond excluding PLK-2 from the nucleoplam by promoting CHK-2 activity. We suggest that in addition to preventing inappropriate localization of PLKs, HAL-2/HAL-3 also antagonizes the activity of PLKs even when they are mis-localized, *e.g.* by keeping them from phosphorylating inappropriate targets and/or by promoting target dephosphorylation. In this scenario, restricting PLK activity to spatially and temporarily appropriate targets would be the combined result of both positive targeting influences that promote association with appropriate substrates at specific subcellular sites, and antagonistic forces that inhibit/counterbalance activity of PLKs toward spatially- or temporally-inappropriate substrates.

### Rethinking the neglected nucleoplasm

Most research on the mechanisms underlying chromosome movement, homolog pairing, SC assembly and recombination during meiosis has quite reasonably focused primarily on proteins or structures interacting directly with the chromosomes themselves, or indirectly with the chromosomes through NE-spanning protein complexes. Here, we have instead focused on the composition and roles of a protein complex that localizes predominantly within the nucleoplasm yet has a profound impact on the behavior and organization of the chromosomes and chromosome-associated structures. The essential role of the HAL-2/HAL-3 complex in coordinating the chromosomal events of meiotic prophase underscores the fact the nuclear environment in which these events occur can have a major influence on their success. It makes sense that this should be the case, as all of the components of the chromosomes and associated structures must traverse the nucleoplasm to reach their eventual subnuclear destinations. Moreover, movement of the chromosomes occurs within the context of the nucleoplasm, so factors that affect the viscoelastic properties of the nucleoplasm would be expected to influence the timing or efficiency of homolog pairing (NABESHIMA 2012).

While misregulation of PLK localization and activity is one major consequence of loss of the HAL-2/HAL-3 complex, our data do not allow us to discern conclusively whether mis-localization/inappropriate activity of PLKs is fully responsible for all of the meiotic defects observed in *hal-2* and *hal-3* mutants. Indeed, we previously found that localization of PLK-2 to PCs can be transiently and partially restored when SYP proteins are eliminated in a *hal-2* mutant background, indicating that aberrant behavior of SYP proteins can contribute to impairment of chromosome movement and homolog pairing (ZHANG *et al.* 2012). Based on our current finding that tripartite SC structure is partially restored by elimination of PLK-2 in the *hal-2* mutant background, we infer that at least some of this “bad behavior” of SYP proteins may be attributable to rogue phosphorylation of chromosome axis and/or SC-CR components by nucleoplasmic PLK activity. However, it is probable that additional meiotic machinery components and/or processes may also be perturbed by an altered state of the nucleoplasm in *hal* mutants. Further, we think it likely that the contributions of nucleoplasmic components in general have been underestimated. There is a growing recognition that proteins localized predominantly in the nucleoplasm can have a profound impact on chromosome structure (e.g. WAPL-1; (CRAWLEY *et al.* 2016)), that many chromosome-associated proteins also have substantial nucleoplasmic pools (e.g. (SILVA *et al.* 2014; JANISIW *et al.* 2018; LI *et al.* 2018; WOGLAR AND VILLENEUVE 2018)), and that dynamic exchange can occur between protein pools in different subnuclear compartments (e.g. (PATTABIRAMAN *et al.* 2017)). Thus, we anticipate that future research will uncover roles for additional nucleoplasmic factors in promoting a successful outcome of the meiotic program.

## ACKNOWLEDGEMENTS

We thank the Caenorhabditis Genetics Center (funded by NIH Office of Research Infrastructure Programs P40 OD010440) for strains, C. Frokjaer-Jensen for the plasmid encoding the germline optimized GFP and N. Bhalla, A. Dernburg, V. Jantsch, E. Kipreos, M. Zetka and R. Lin for antibodies. This work was supported by MRC grant MC-A652-5PY60 to EM-P and by American Cancer Society Research Professor Award (RP-15-209-01-DDC) and NIH grants R01GM53804 and R35GM126964 to AMV. The project described was also supported, in part, by Award Number 1S10OD01227601 from the National Center for Research Resources (NCRR) to the Stanford Cell Science Imaging Facility. Its contents are solely the responsibility of the authors and do not necessarily represent the official views of the NCRR or the National Institutes of Health.

## Supporting Information

**Figure S1:**
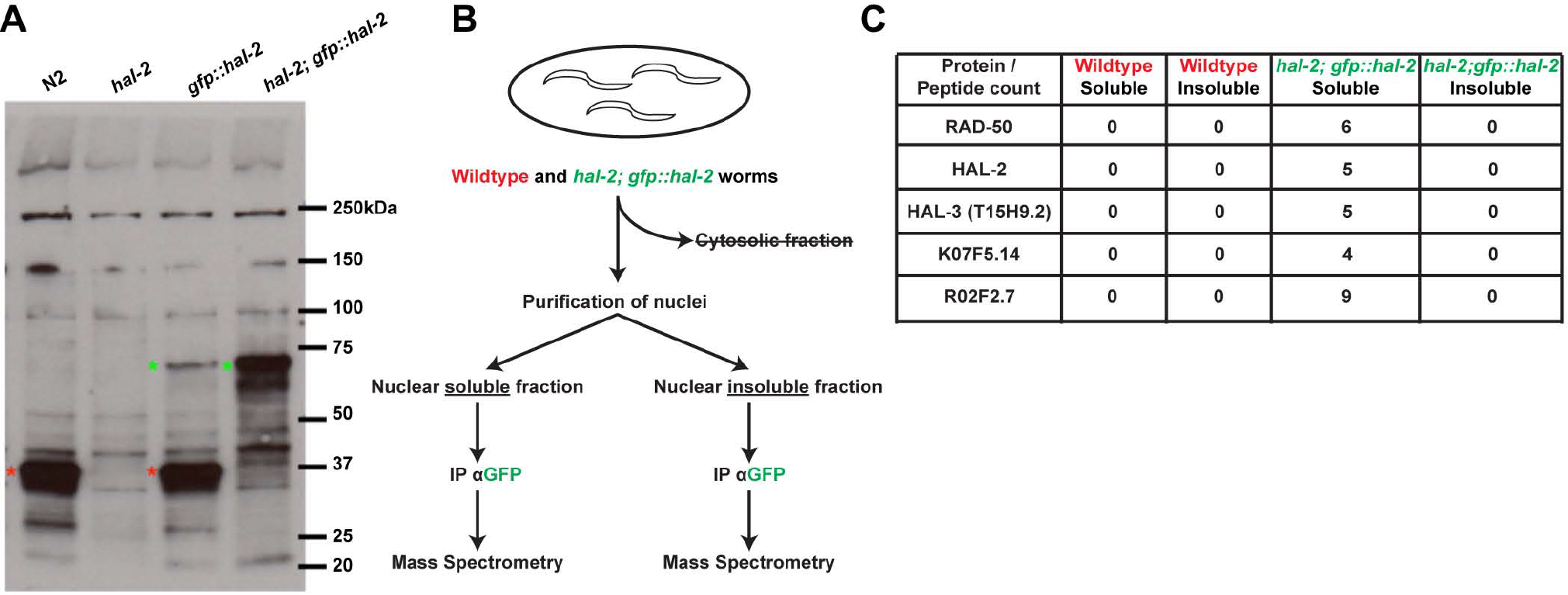
A proteomic based approach to identify binding partners of HAL-2. **A)** Detection of HAL-2 protein in whole worm lysates of the indicated genotype. HAL-2 expressed from the endogenous locus (36kDa, red asterisks) and GFP::HAL-2 expressed from an integrated transgene (62kDa, green asterisks) were detected by Western blot analysis performed using rabbit anti-HAL-2 antibody ((ZHANG et al. 2012) at 1:10,000 dilution). As the level of GFP::HAL-2 is higher in the *hal-2(me79)* homozygous mutant background (lane 4) than in otherwise wild-type animals (lane 3) the *hal-2; gfp::hal-2* strain was therefore used for our proteomics strategy. **B)** Schematic representation of the proteomics strategy used to identify HAL-2 partners (see Materials and Methods). HAL-2 peptides were detected only in the nuclear soluble fraction derived from worms expressing GFP::HAL-2, consistent with Western blot analyses of fractionated extracts (SILVA et al. 2014). 20 additional proteins exhibited this same profile, including RAD-50, which has previously-demonstrated roles in meiotic recombination. Additional criteria were applied to define a set of candidate genes among the remaining 19 to test for roles in regulating meiotic prophase events. Eight genes were excluded because they are located on the X chromosome (which lacks known meiosis genes) and/or encode tubulin isoforms. Two additional genes were excluded based on evidence that they are likely not expressed in the germ line (Wormbase). Six genes were not pursued further based on prior annotation/functional analysis, including three genes encoding proteins (PGL-1, CGH-1 and OMA-1) that are known to localize to P-granules and/or cytoplasm and have previously-described roles in translational regulation. The remaining genes (*T15H9.2, K07F5.14, R02F2.7*) were prioritized as candidates for further study based on strong evidence for germline-enriched expression (REINKE et al. 2000; KIM et al. 2001; HAN et al. 2017), yet little or no prior in-depth analysis of germline function. **C)** Table showing numbers of peptides detected for HAL-2, RAD-50, and the three prioritized candidate proteins (with peptides detected specifically in the nuclear soluble fraction from worms expressing GFP::HAL-2.

**Figure S2:**
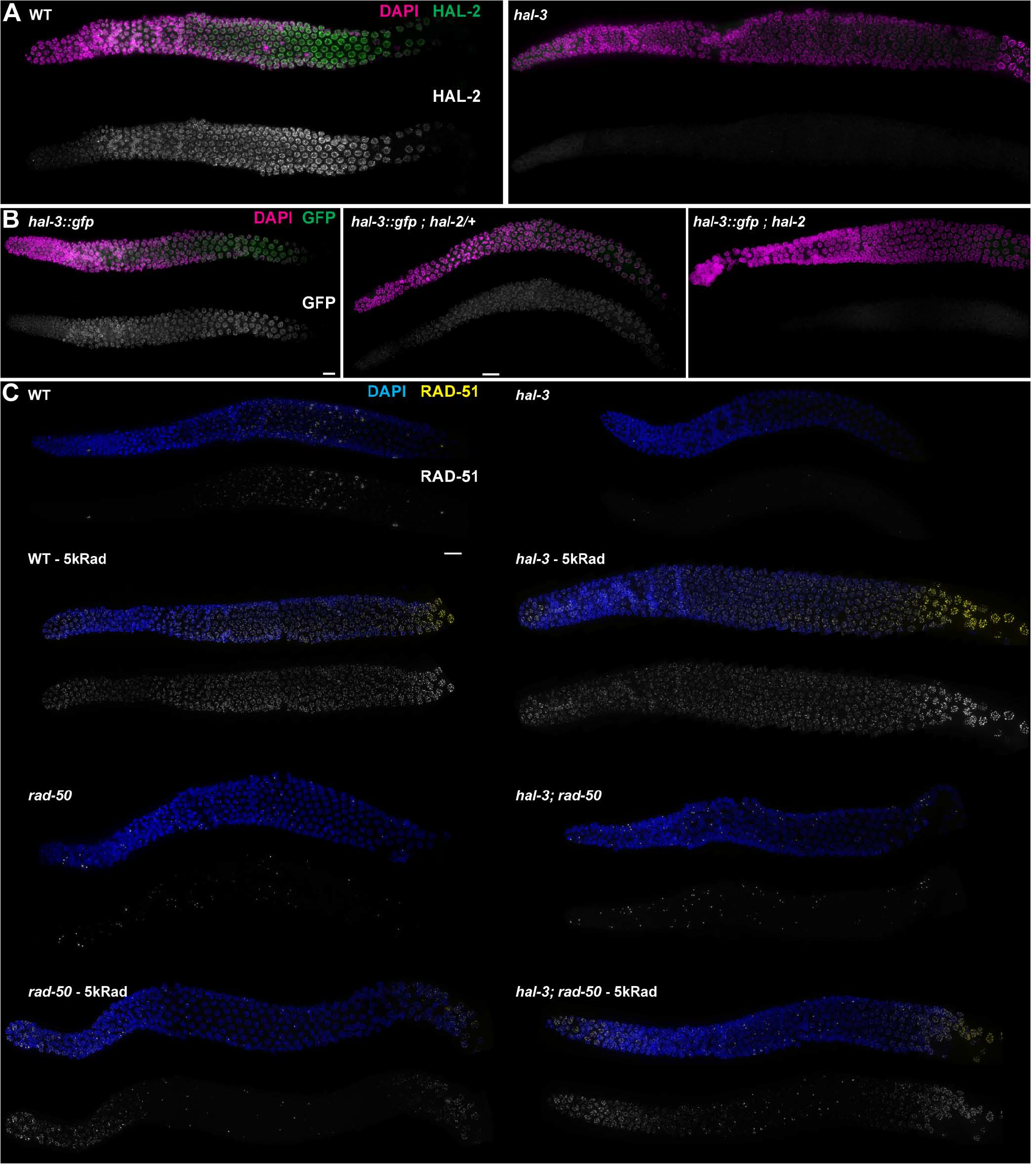
Whole gonad images of samples presented in Figure 1. **A)** Immunodetection of HAL-3::GFP fusion protein in whole mount gonads of worms of the indicated genotypes. Samples are the same as in Figure 1E. Importantly, the fusion protein could be detected in *hal-2*/+ heterozygous worms (middle panel) but not in their *hal-2/hal-2* homozygous mutant siblings (right panel), indicating that transgene silencing is unlikely to account for the lack of signal observed in the *hal-2/hal-2* homozygote. **B)** Immunodetection of HAL-2 in whole mount gonads of worms of the indicated genotypes. Samples are the same as in Figure 1D. C) Immunodetection of RAD-51 in whole mount gonads of worms of the indicated genotypes, either unirradiated or one-hour post exposure to 5kRad of γ-irradiation. Samples are the same as in Figure 1F. Abundant radiation-induced RAD-51 foci are detected in nuclei throughout the entire depicted portions of the wild type and the *hal-3* single mutant germ lines, whereas radiation-induced foci are limited to the premeiotic and late pachytene regions in the *rad-50* and *hal-3; rad-50* germ lines. In all panels, images are maximum intensity projections of the top layer of nuclei in the imaged gonads; scale bars represent 20μm.

**Figure S3:**
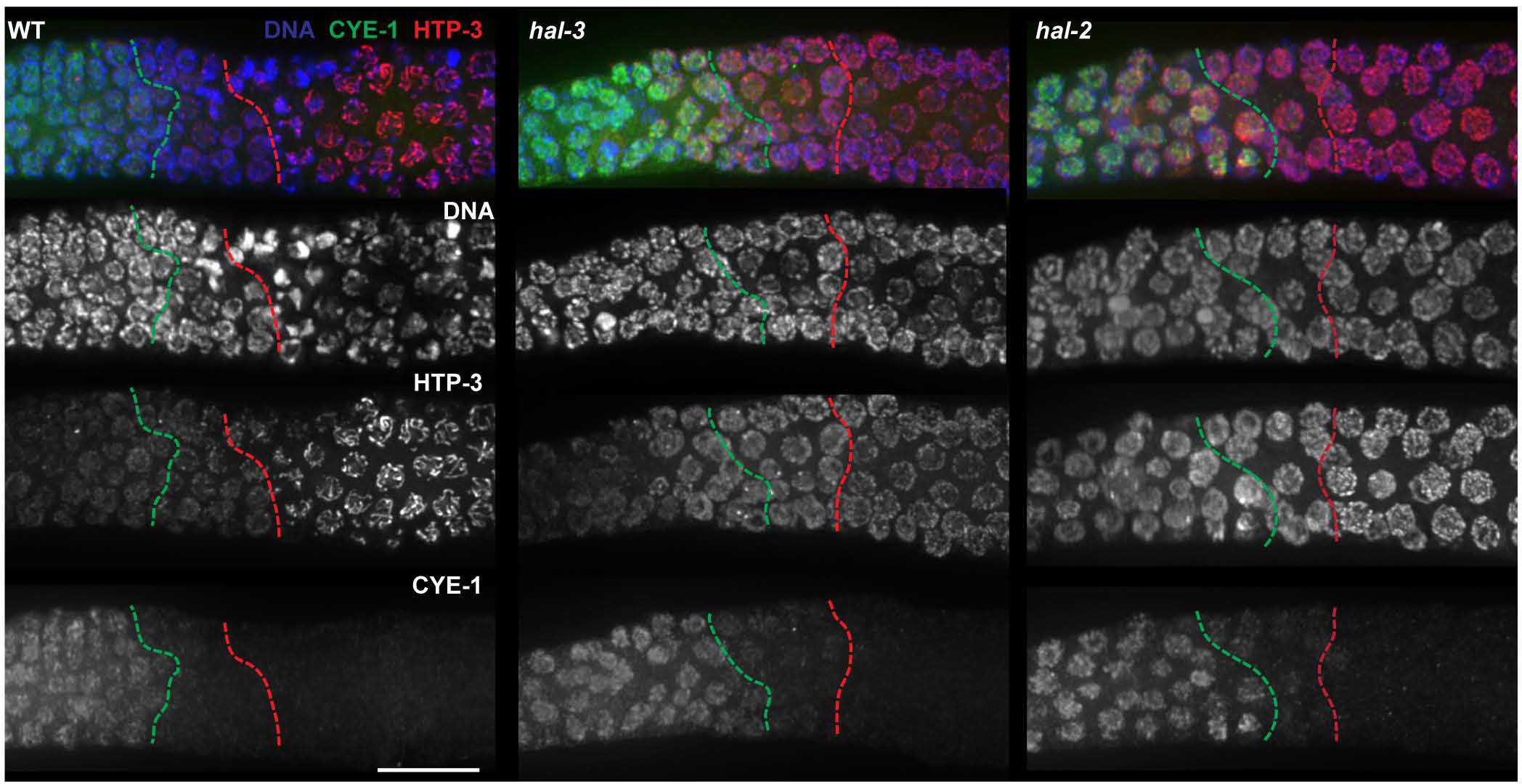
Mitosis to meiosis transition is normal in *hal-2* and *hal-3* mutants. Zoomed-in images of the meiotic entry region of the germ lines depicted in Figure 2A, providing merged images of mitotic cell cycle marker Cyclin E1 (CYE-1), chromosome axis marker HTP-3, and DAPI-stained DNA, together with single channel images. The green dashed lines represent the last row of nuclei with high CYE-1 signal; the red dashed lines represent the first row of nuclei with fully assembled chromosome axes. Scale bar represents 20μm.

**Figure S4:**
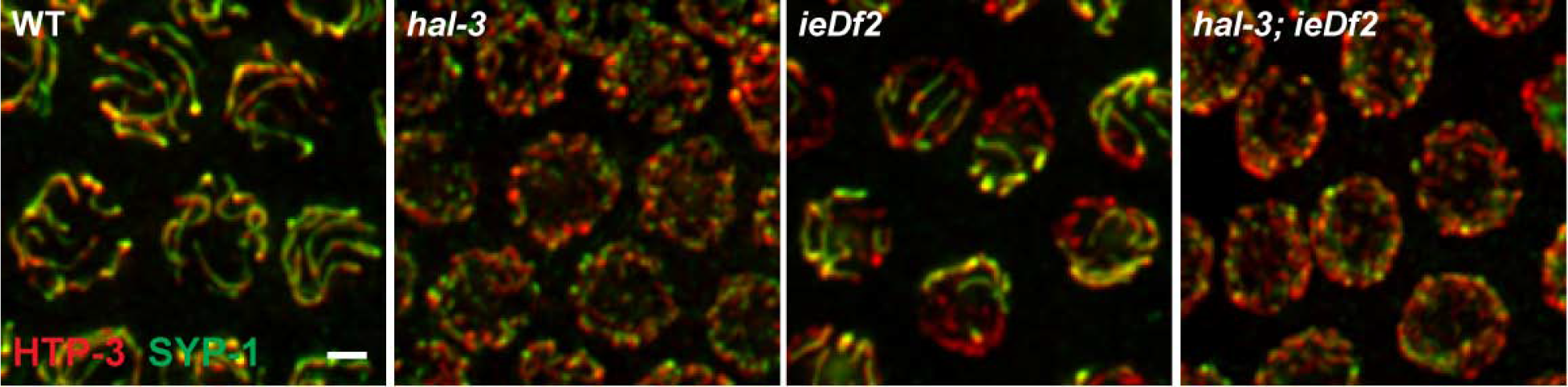
Lack of PC-binding protein or *hal* lead to different defects in synapsis. Nuclei from the mid-pachytene region immunostained for HTP-3 (red) and SYP-1 (green), illustrating the differences in appearance of nuclei with normal full synapsis (WT), delayed foldback synapsis on a subset of chromosomes (*ieDf2*), or loading of SYP-1 along the lengths of individual unpaired axes (*hal-3 and hal-3; ieDf2*). Scale bar represents 2μm.

**Figure S5:**
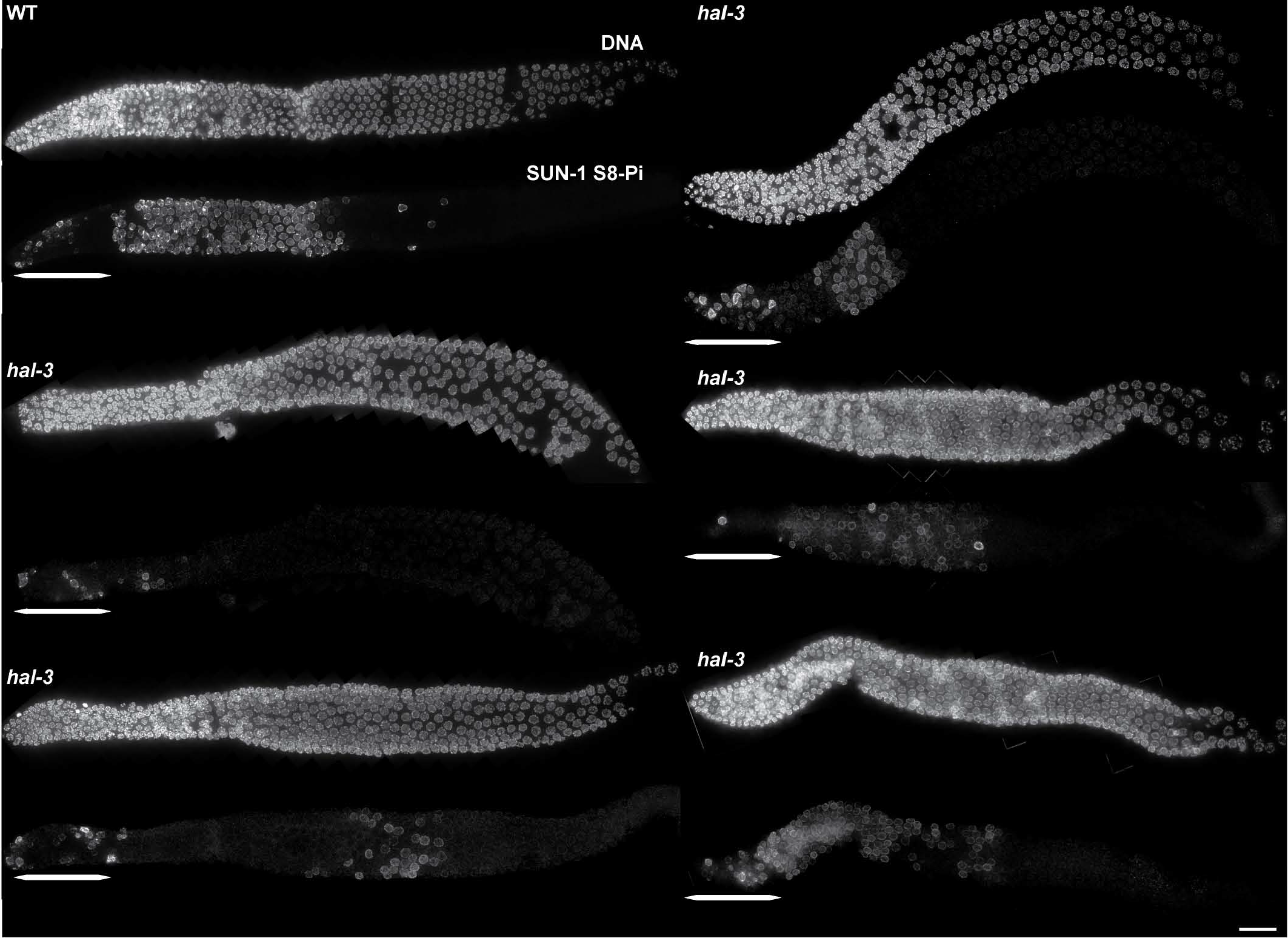
Variability in the number and position of meiotic prophase nuclei positive for SUN-1 S8-Pi in *hal-3* mutant gonads. Immunodetection of the nuclear envelope protein SUN-1 phosphorylated on serine 8 (SUN-1 S8-Pi), a marker of CHK-2 activity in early meiotic prophase. The wild type germ line illustrates the normal pattern, where SUN-1 S8Pi is robustly detected in meiotic nuclei from the onset of meiotic prophase through the end of early pachytene stage. The five *hal-3* samples were chosen to illustrate the high variability observed in *hal-3* mutant germ lines, both in the number of nuclei positive for this marker of early prophase and in the distribution of such nuclei across the meiotic prophase region. In the premeiotic region of the gonad, highlighted by a bracket, SUN-1 S8-Pi is detected in nuclei undergoing mitosis but at this stage, the modification is independent of CHK-2 (PENKNER et al. 2009).

**Figure S6:**
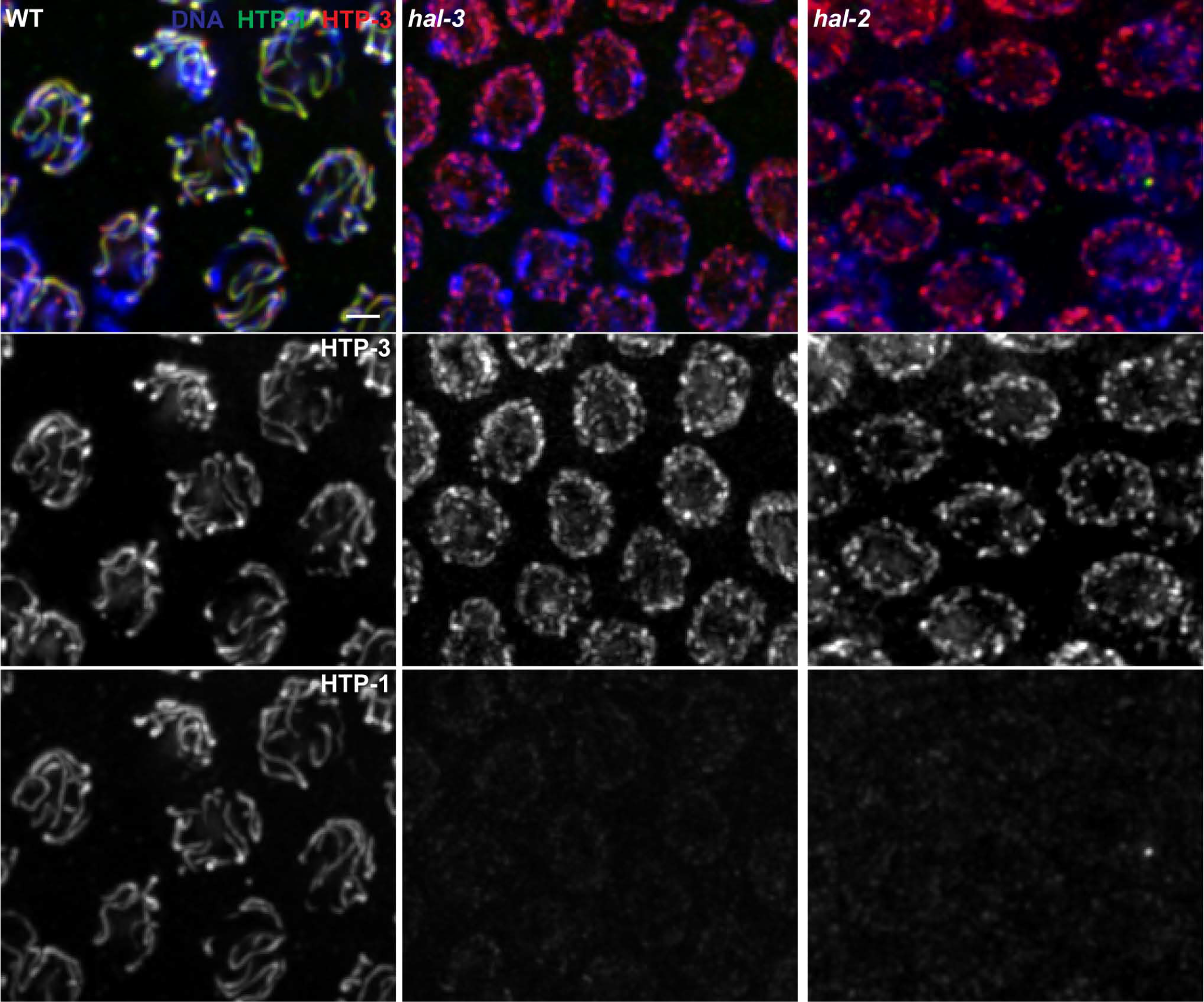
Defective chromosomal association of axis protein HTP-1 in *hal* mutants. Immunodetection of meiotic chromosome axis proteins HTP-3 (red) and HTP-1 (green), and DNA counterstained with DAPI (blue) in early pachytene nuclei of the indicated genotype indicates that, as previously described for *hal-2* (ZHANG et al. 2012), levels of axis-associated HTP-1 are strongly reduced in *hal-3* mutant meiocytes.

